# Resolution of multi-receiver trade-offs in wall lizard colouration

**DOI:** 10.64898/2026.08.25.746845

**Authors:** Lekshmi B Sreelatha, Javier Abalos, Prem Aguilar, Alexandra M. Tyers, Ossi Nokelainen, Zbyszek Boratyński, Miguel Angel Carretero

## Abstract

Animal colouration evolves under multiple, often conflicting, selective pressures. Conspicuous, non-aposematic colour-patterns that enhance conspecific communication may simultaneously increase detectability by predators. Such trade-offs can be resolved by optimising colour-patterns to match the perceptual abilities of different receivers. We tested whether dorsal colour-patterns of the Lusitanian wall lizard (*Podarcis lusitanicus*) are optimised for ecologically relevant receivers across relevant viewing distances, while accounting for the visual acuity of conspecifics and predators. Conspecifics and snakes detected chromatic information at shorter distances, whereas achromatic and luminance information were detected at longer distances. Birds showed a uniform decline in detectability across the colour-pattern components with increasing viewing distance. Larger males retained high chromatic detectability across all receivers despite the general distance-related decline, whereas females and smaller individuals exhibited less salient colour patterns, consistent with predator avoidance strategy. Our results show that lizards resolve the trade-off between conspecific communication and predator detection through distance-dependent colour-pattern perceptibility across receivers. This resolution breaks down in large males, for whom the benefits of salient chromatic patterns for intraspecific communication may outweigh increased detectability to predators.

## Introduction

Animal colouration mediates multiple functions, from physiological roles such as thermoregulation, photoprotection and immune defence (Hegna et al., 2013; Goldenberg et al., 2024; Sacchi et al., 2007), to communication with conspecifics and predators (Cuthill et al., 2017). Colour patterns are central for visual signalling, shaping interactions such as mate choice, intrasexual competition and predator avoidance (Boratyński et al., 2017; Caro & Allen, 2017; Nokelainen et al., 2024; Vilela et al., 2017; Nokelainen et al., 2017). Consequently, animal colour patterns are shaped by multiple, often conflicting, selective pressures (Cuthill et al., 2017), often reflecting a balance between the benefits of socio-sexual signalling and the risk of predation (Amdekar & Thaker, 2019; Storniolo et al., 2021). Conspicuous colour patterns can enhance mate attraction and intimidation of rivals but also render individuals more detectable to predators whose visual systems are adapted to exploit such signals (Lebas & Marshall, 2000; Stuart-Fox et al., 2003; Pérez i de Lanuza et al., 2013). Given substantial variation among predator visual systems, selection on colour patterns may reflect the perceptual abilities of the dominant visual predators (Endler, 1992; Hart, 2002; Nokelainen et al., 2020; Sillman et al., 1997). The strong and opposing selective pressures are likely to resolve such trade-offs, whereby colour patterns are optimised for efficient intraspecific communication, while minimising detection by predators (Endler, 1980; Endler, 1978; Marshall & Stevens, 2014).

Reptiles, particularly lizards, provide excellent models for examining the evolution of animal colouration, as colour patterns are expected to be simultaneously shaped by sexual and natural selection, influencing communication, thermoregulation and camouflage (Clusella-Trullas et al., 2007; Sreelatha et al., 2021; Storniolo et al., 2021; Stuart-Fox & Ord, 2004). Colour-pattern variation driven by sexual selection may be limited by ecological and behavioural adaptations to predation avoidance and habitat use, where conspicuousness can be offset by compensatory anti-predator strategies (Ortega et al., 2014, 2015; Stuart-Fox & Ord, 2004; Vilela et al., 2017). Empirical evidence supports this interplay: for example, male Italian wall lizards (*Podarcis siculus*) display bright dorsal colouration potentially relevant for communication, whereas females show more cryptic patterns linked to camouflage (Storniolo et al., 2021). In some populations of *Podarcis erhardii*, males partition colour signals across body regions with differing visibility to receivers, dorsal surfaces exposed to aerial predators are better matched to the background, while ventrolateral flanks visible to conspecifics are more conspicuous, whereas females show uniformly cryptic colouration across the body (Marshall & Stevens, 2014). Individual body size may modulate these dynamics, as it affects the spatial scale (amount of information) over which colour-pattern components are projected to receivers, potentially amplifying receiver-dependent trade-off between communication and crypsis (Echeverri et al., 2021).

We hypothesise that the dorsal colour patterns of Lusitanian wall lizard, *Podarcis lusitanicus,* reflect a resolved trade-off, optimised for conspecific communication while maximising crypsis from predators. Although dorsal colour patterns in lacertid lizards have received less attention than ventral or lateral colour patches as targets of selection (Pérez i de Lanuza et al., 2014; Marshall & Stevens, 2014), they may nonetheless be shaped by sex-specific selective pressures. Specifically, males and females may differ in dorsal colour-pattern detectability if they experience different selection arising from conspecific interactions and predator avoidance (Marshall & Stevens, 2014; Storniolo et al., 2021). *P. lusitanicus* lizards exhibit pronounced dorsal colour-pattern variation across the diverse habitats of north-western Iberia (Sreelatha et al., 2025), making it well suited for testing receiver-dependent colour-pattern trade-offs. To test whether this trade-off is resolved by exploiting receiver-specific visual perception, we modelled the dorsal colour patterns of *P. lusitanicus* as perceived by conspecifics and their principal predators, snakes and birds (Marshall & Stevens, 2014; Martin et al., 2015; Pérez i de Lanuza & Font, 2015). We specifically tested: (1) whether colour patterns balance detectability by conspecifics and predators across viewing distances; (2) Whether detectability of sex-specific information available for intraspecific communication declines with viewing distance for all receivers but more strongly for predators; and (3) whether large individuals lose the trade-off resolution, owing to the greater amount of information their larger bodies project. We also expect that colour-patterns as perceived by the conspecifics are more evolutionarily conserved, in contrast to the predator-perceived components, as predator-mediated selection may vary spatially with differences in local predator community composition.

## Methods

### Study species

The Lusitanian wall lizard, *Podarcis lusitanicus* Geniez, Sá-Sousa, Guillaume, Cluchier & Crochet, 2014, is a small lacertid lizard (∼1.5-5g), endemic to the northwestern Iberian Peninsula. It is a diurnal, mostly saxicolous, using rock (granite and schist) surfaces for thermoregulation, refuge, and foraging, to a larger extent than other *Podarcis* species (Gomes et al., 2016). Using this system, previous research has shown that environmental variables shape dorsal lightness in *P. lusitanicus* (Sreelatha et al., 2025). Here, we analysed a dorsal image dataset comprising 470 adult *P. lusitanicus* (267 males, 203 females), captured by noosing during their daily activity period in the reproductive season (Carretero et al., 2022), between April and July of 2022 (18 populations in Portugal) and 2023 (three populations in Galicia, Spain; Figure. S1). For each individual, we took multispectral images (detailed below) and measured snout-vent length (SVL: body size hereafter; to the nearest 0.01 mm using an electronic calliper; mean ± SE = 51.55 ± 0.30 mm) and body mass (to the nearest 0.01 g using a portable digital scale; 2.66 ± 0.05 g). Lizards were released afterwards at their location of capture within the same day.

### Photography and image calibration

Visible (VIS) and ultraviolet (UV) spectrum photographs were taken using a customised full spectrum Samsung NX1000 camera equipped with minimum light absorption Novoflex Noflexar 35 mm lens, in daylight under a photographic umbrella, as in Sreelatha et al. (2025). Multispectral images were produced by merging pairs of VIS and UV camera RAW images (Samsung SRW files), selecting only those with the highest photographic quality to reduce analytical error. All image analyses were performed in ImageJ (Abràmoff et al., 2004) using the Multispectral Image Calibration and Analysis Toolbox (micaToolbox) v.2.3 (Troscianko & Stevens, 2015). Visible and UV photographs were aligned and normalized using a 50% reflectance standard to standardise radiance and lighting conditions, and images were rescaled to a 30 mm scale bar (Stevens et al., 2007). Using the polygon tool in ImageJ, the dorsal body region, excluding the head, limbs and tail, was selected as the region of interest (ROI) for analysis (for details see: Sreelatha et al. 2025).

### Quantitative colour pattern analysis (QCPA) and vision modelling

To model the appearance of lizard dorsum for receiver vision systems, we used the micaToolbox ImageJ plugin and the quantitative colour pattern analysis pipeline producing cone-catch images (Batch QCPA, v1.3; van den Berg et al., 2020, 2024). The QCPA framework combines calibrated digital photography, visual modelling, and spatio-chromatic colour-pattern analyses to simulate how visual scenes are perceived by different observers, accounting for their photoreceptor sensitivities, receptor noise, light environment, spatial acuity, and viewing distance. QCPA incorporates receptor noise-limited (RNL) clustering for image segmentation and RNL-ranked filtering for visual acuity modelling of different observers, along with several complementary analyses of colour pattern (Figure. 1a; van den Berg et al., 2020) such as colour adjacency (Endler, 2012), visual contrast (Endler & Mielke, 2005), boundary strength (Endler et al., 2018), and local edge intensity analysis, LEIA (van den Berg et al., 2020).

**Figure 1.**
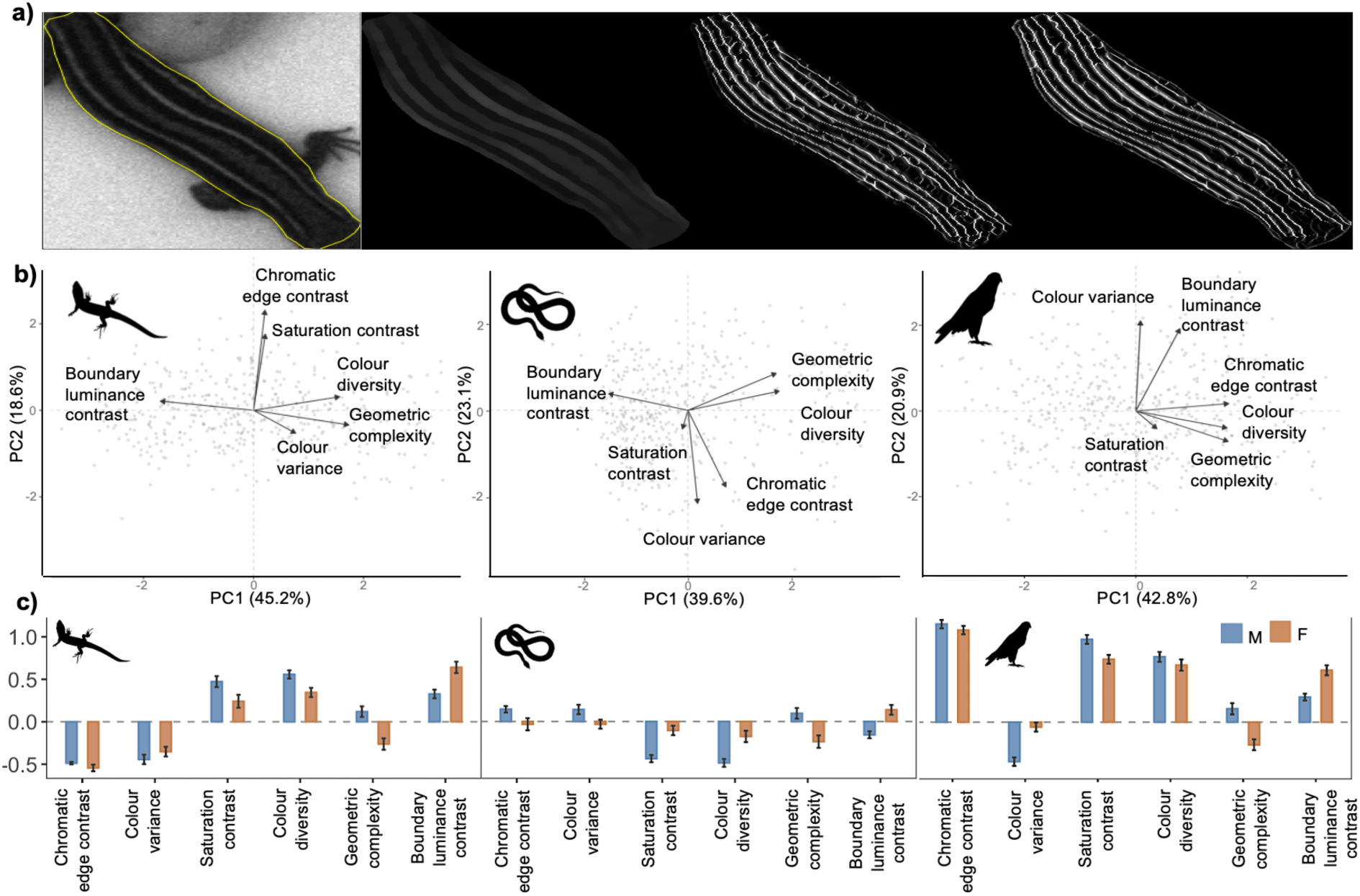
(a) Representative images showing different stages from the Quantitative Colour Pattern Analysis (QCPA). From left to right: region of interest selected (using ImageJ) for QCPA from the dorsum of *Podarcis lusitanicus* wall lizards; acuity corrected (for bird vision at 1.5m) image eliminating the details the specific vision system cannot resolve; chromatic edges from local edge intensity analysis (LEIA) after reconstructing the edges with RNL (receptor noise limited) clustering; and luminance edges from LEIA after reconstructing the edges with RNL clustering. (b) Principal component analysis (PCA) biplots of dorsal colour-pattern components as perceived through conspecific lizard, snake, and avian predator vision systems at 1.5 m viewing distance. Grey points represent individual lizards in multivariate colour-pattern space. Arrows show the loadings of six colour-pattern variables on PC1 and PC2, and the percentage of total variance explained by each axis is given in parentheses. (c) Standardised values (± S.E) of six colour-pattern variables (chromatic edge contrast, colour variance, saturation contrast, colour diversity, geometric complexity, boundary luminance contrast) for males (blue) and females (orange), shown separately for lizard, snake, and bird vision systems. All silhouettes were obtained from PhyloPic.org. The conspecific lizard silhouette is attributed to Titouan Montessui (licensed under CC BY 4.0, https://creativecommons.org/licenses/by/4.0/). The violet-sensitive bird and snake silhouettes were sourced from openly licensed images.

We modelled the dorsal ROI of *P. lusitanicus* under three visual systems representing potential predators (birds and snakes) and conspecific lizards, across relevant viewing distances, outlined below. The main risk of predation on this lizard species originates from raptors (e.g., *Buteo buteo*, *Falco tinnunculus*), corvids (*Corvus* spp.) (Marshall & Stevens, 2014), and snakes (Pérez i de Lanuza & Font, 2015) such as *Coronella austriaca* (Carretero et al., 2022). Cone-catch models were generated using the spectral sensitivities available in micaToolbox for the tetrachromatic peafowl (*Pavo cristatus*) vision system (Hart, 2002), as a proxy of the violet-sensitive (VS) vision of the major bird predators. For the snake vision system, as no data are currently available for *C. austriaca*, we used the trichromatic spectral sensitivities of another colubrid, *Thamnophis sirtalis* (Goedert et al., 2021; Sillman et al., 1997). For the lizard conspecific model, we used the tetrachromatic spectral sensitivities of *Podarcis muralis*, a congener with comparable visual ecology (Abalos et al., 2025, Martin et al., 2015).

The ability to resolve colour pattern details depends on the observer’s spatial visual acuity and the viewing distance (Endler & Mappes, 2017; Salisbury & Peters, 2025). We set the spatial acuity (Table S1) to 2.05 cycles per degree (cpd) for the conspecific lizard model (Kawamoto et al., 2025), 40 cpd for the avian predator model (Hirsch, 1982), and 4 cpd for the snake predator model (Baker et al., 2007). We modelled the dorsal ROI under each vision system at three viewing distances (short, medium, and long; see below) Distance values differed among observers because they were defined according to ecologically relevant viewing ranges and species-specific spatial acuity (Table S1, Baker et al., 2007; Hirsch, 1982; Kawamoto et al., 2025; Simões et al., 2016). Preliminary analyses across a wider distance range showed that, beyond the selected thresholds, the dorsal ROI no longer retained sufficient spatial resolution for quantitative colour pattern assessment. We therefore restricted subsequent analyses to three ecologically relevant distances that maintained adequate data quality (Salisbury & Peters, 2025). Specifically, we modelled violet-sensitive bird vision (hereafter bird vision) at 1.5, 3, and 4 m; snake vision at 1, 1.5, and 3 m; and lizard vision at 0.5, 1.5 and 2 m, corresponding to short, medium, and long viewing distances, respectively. Receptor sensitivities (λmax), Weber fractions, and spatial acuity values are given in Table S1.

The multispectral images were run through the batch QCPA framework (van den Berg et al., 2020, 2024). Batch processing allowed the application of spatial acuity corrections across multiple viewing distances within a single pipeline. Each image was processed in three independent runs, one for each vision model. In summary, the workflow (Figure 1a) included: (i) Gaussian acuity correction based on the combination of viewing distance and the observer’s spatial acuity, removing pattern elements that the visual system cannot resolve (van den Berg et al., 2020); (ii) receptor noise-limited (RNL) filtering and clustering, producing an image segmented according to the perceptual discrimination thresholds of the viewer (van den Berg et al., 2020); and (iii) automated colour pattern analyses, including Colour Adjacency Analysis (CAA, Endler, 2012), Visual Contrast Analysis (VCA, Endler & Mielke, 2005), Boundary Strength Analysis (BSA, Endler et al., 2018), particle analysis, and Local Edge Intensity Analysis (LEIA, van den Berg et al., 2020), which assess the contrast, complexity and heterogeneity of the colour patterns. These analyses generate a comprehensive set of colour-pattern variables, quantifying chromatic and achromatic properties (such as colour and brightness), pattern geometry, and their integration into spatio-chromatic measures of pattern contrast (van den Berg et al., 2020). From the 157 colour-pattern variables generated by the QCPA pipeline, we first filtered out redundant parameters (i.e. highly correlated: r ≥ 0.7; Figure. S2 for details). From the remaining variables, we selected six that collectively capture the variation in the colour pattern of our study system: (1) chromatic edge contrast, (2) colour variance, (3) saturation contrast, (4) colour diversity, (5) geometric complexity, and (6) boundary luminance contrast. Variables 1-3 quantify the chromatic edge strength and saliency of chromatic information available to receivers, while variables 4 -6 capture the colour/luminance classes within the spatial patterning, achromatic pattern structure and luminance boundaries of the colour pattern.

### Statistical analysis

We examined the mean, standard deviation (SD), skewness and kurtosis of each colour-pattern variable within each vision system to assess trends and identify potential extreme values in data (Table S2). Observations producing heavy tails, falling outside the 2.5th and 97.5th percentile thresholds of the data distribution, were excluded as outliers. We used multivariate analysis of variance (MANOVA) as a global test, including population, sex, body size and their interactions as explanatory variables, to evaluate whether colour-pattern components differed among populations, separately for each visual system.

We analysed how viewing distance influences detectability of colour-pattern components of males and females as ‘perceived’ under three different vision models (conspecific lizards and bird and snake predators), using multivariate Bayesian mixed models, implemented in brms (v. 2.22.0; Bürkner, 2017) in R (v.4.4.0; R Core Team. R Foundation for Statistical Computing, 2024). Six models were fit (three visual systems × two sexes; sex-specific models avoiding confounding sex effects), each of which simultaneously estimated six standardised QCPA-derived colour-pattern variables (chromatic edge contrast, colour variance, geometric complexity, colour diversity, saturation contrast, boundary luminance contrast) with identical fixed effects. Each colour-pattern variable (log transformed, scaled and centred) was modelled as a separate Gaussian response with viewing distance (categorical) as a fixed effect and body size (log transformed, scaled and centred) as a covariate to control for its potential influence on colour-pattern components (Sreelatha et al., 2025). To account for repeated measures of individuals for different distances and sampling structure, we included random intercepts for individual lizards nested within sampling location. We ran four chains with 15,000 iterations each (1,000 warmup and thinning of 10, yielding 5600 post warm up samples) using weakly informative priors. To account for shared unexplained variance and inherent covariance structure of colour-pattern variables, we specified set_rescor = TRUE. Convergence and sampling quality of multivariate models were assessed using Rhat (all ≤ 1.01) values, effective sample sizes (> 4105), and posterior predictive checks. Model performance was assessed using Pareto-smoothed importance sampling leave-one-out cross-validation (PSIS-LOO; Vehtari et al., 2016). Pareto k diagnostics and LOO plots were examined to identify potentially influential observations (k>0.7). Flagged cases were retained because they did not affect posterior estimates or model interpretation.

To test the effect of different predictors on the colouration of males and female (models for both sexes were identically parametrised; same predictors and settings, and similar priors), we calculated: (1) the percentage overlap of their posterior distributions (overlap %) and credible intervals (overlap CI %); lower values indicate more divergent responses between males and females), and (2) the difference between male and female effect sizes. For these, we computed posterior distributions of the difference (males – females) for each variable. We considered that the colouration of males and females responded credibly differently to a predictor if the 95% highest density interval (HDI, the range containing 95% of the most probable difference values) did not include zero (Kruschke, 2012). Although sample sizes differed slightly between sexes (male: female ratio ∼1.3:1, Table S2), both groups had sufficient observations (>500, reflecting three distance-level measurements per individual) for robust posterior estimation. For predictor effects with low overlap between sexes, we evaluated the probability that male (M) effects exceeded female (F) effects, by calculating probability of direction (pd _(M>F)_, defined as the proportion of the posterior distribution supporting a particular direction, Makowski et al., 2019). Values approaching 1 indicate high certainty that males show stronger effects, while values approaching 0 indicate females show stronger effects.

To test whether the colour patterns as perceived by the conspecifics are more evolutionarily conserved in contrast to the predator perceived components, we tested for a phylogenetic signal (population genetic structure) by fitting population-level means of colour-pattern components measured at 1.5 m distance across all visual systems, using generalized linear mixed models in a Markov chain Monte Carlo framework (MCMCglmm v. 2.36; Hadfield, 2010; Stone et al., 2011). Within each vision system, separate univariate models were fitted for each colour pattern variable (200,000 iterations, 8,000 burn-in, thinning = 100) with an intercept only fixed effect structure, with population genetic distances derived from pairwise *F_ST_* values from Rato et al. (2024), estimated for the same populations and individuals as a random effect.

## Results

### Variation in phenotype

At a common viewing distance of 1.5 m, colour-pattern components varied broadly across individuals (Figure 1b), with males showing consistently higher mean chromatic trait values and females showing higher mean boundary luminance contrast across all three vision systems (Table S2, Figure 1c). A significant effect of population on colour-pattern components was detected across all visual systems, and population was therefore included as a factor in all subsequent tests (Table S3). Across populations, in general, males were significantly larger than females: post hoc Tukey test; estimate (female − male) ± SE = −0.46 ± 0.10, df = 422, Tukey-adjusted p < 0.001.

### Colour patterns detectable to conspecifics are less optimal for predators over distances

Our results broadly follow our hypothesis; dorsal colour-pattern components of *P. lusitanicus* showed distance-dependent variation in detectability. But the nature of this variation differed between chromatic and achromatic traits and across vision systems (Figure 2, Figure S3, Table S4). Chromatic traits (chromatic edge contrast, colour variance and saturation contrast) generally declined with increasing distance under conspecific lizard and snake vision, while pattern structure and achromatic edge definition (geometric complexity, colour diversity, and boundary luminance contrast) became more prominent at greater distances under the same vision systems. In contrast, under bird vision chromatic traits remained comparatively stable with distance despite fading chromatic edges, while achromatic edge definition declined with distance (Figure 2, Figure S3, Table S4).

**Figure 2.**
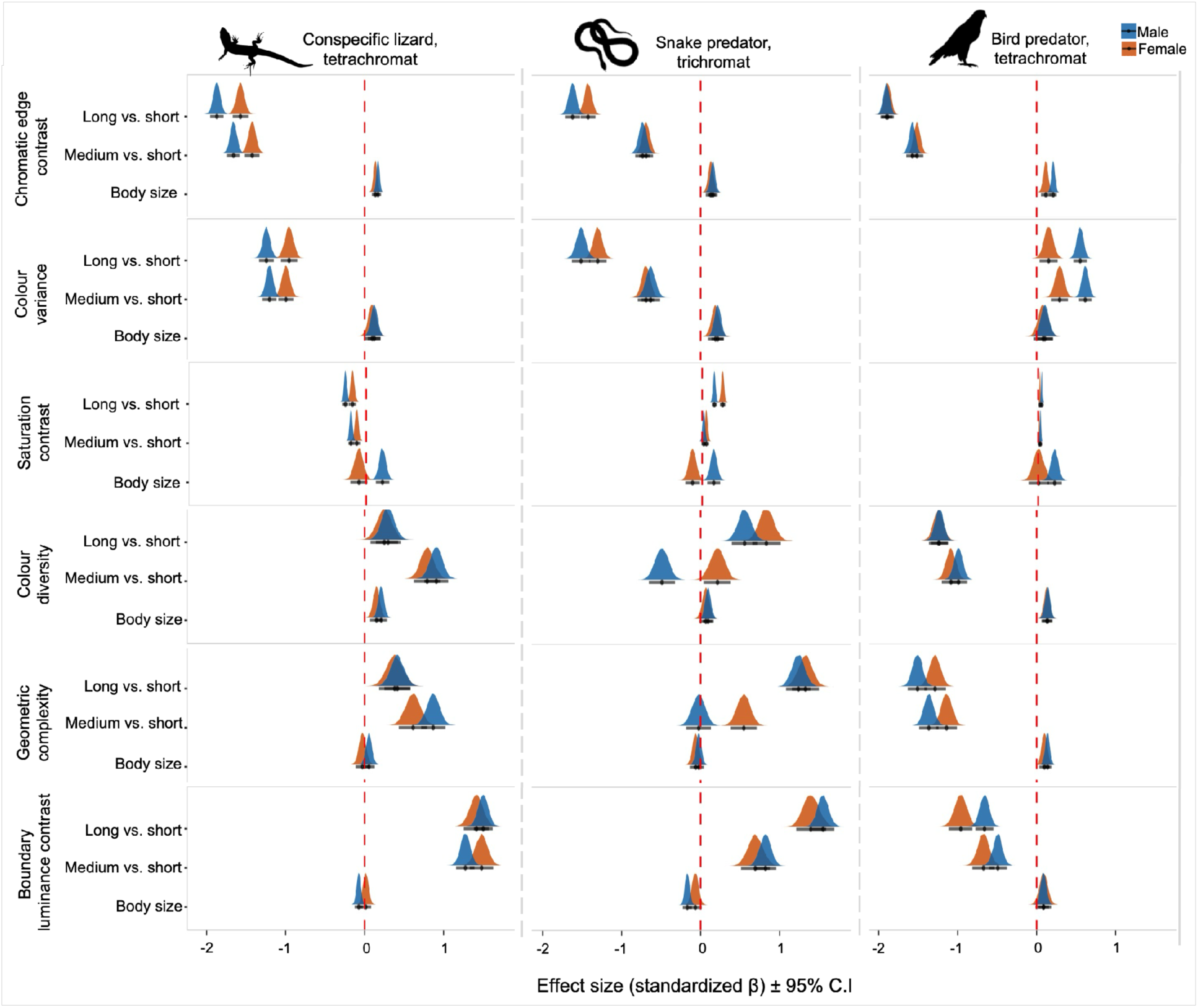
Effects of viewing distance (long vs. short, medium vs. short) and body size on the dorsal colour pattern components of *Podarcis lusitanicus* lizards across different observer vision systems; left panel: lizard vision, middle panel: violet sensitive bird vision, right panel: snake vision. For each vision system, colour pattern descriptors derived from the quantitative colour-pattern analysis (QCPA) were modelled using Bayesian multivariate mixed models, with viewing distance and body size as predictors. Males and females were analysed separately. Individual lizards nested in population is added as random factor. Estimates are reported as mean ± 95% credible intervals (CI). The red dashed line indicates zero effect. Viewing distances modelled: 1) conspecific lizard: 0.5m (short), 1.5m (medium), 2m (long); 2) Bird predator: 1.5m (short), 3m (medium), 4m (long); (3) snake predator: 1m (short), 1.5m (medium), 3m (long). All silhouettes were obtained from PhyloPic.org. The conspecific lizard silhouette is attributed to Titouan Montessui (licensed under CC BY 4.0, https://creativecommons.org/licenses/by/4.0/). The violet-sensitive bird and snake silhouettes were sourced from openly licensed images.

### Sexes differ in trade-off resolution through distance-dependent colour-pattern detectability

Our results support this hypothesis: distance-dependent changes in colour-pattern saliency differed between the sexes across conspecific and predator vision systems, indicating that each sex is tuned to a different balance of detectability by intended versus unintended receivers. Posterior estimates of multiple colour-pattern components showed minimal to no overlap between males and females in different vision systems (Table S5). Under conspecific lizard and snake vision, male colour patterns were optimised for close-range saliency; males had greater decline in chromatic saliency with distance than females, with credible differences in chromatic edge contrast, colour variance, and saturation contrast (probability that males have higher values than females: pd _(M>F)_ < 0.01 in all cases; Table S5, Figure S3). Under bird vision, the pattern reversed: females showed greater declines than males in boundary luminance contrast, saturation contrast and colour variance with distance (pd _(M>F)_ > 0.99), while chromatic edge contrast showed no credible sex difference across distance. Geometric complexity showed vision-system-dependent sex differences, with greater declines for females under lizard vision (pd _(M>F)_ = 0.98), but greater declines for males under snake and bird vision (pd _(M>F)_ < 0.01, Table S5).

### Large individuals fail to resolve the trade-off due to the amount of information projected

Our results only partially support this hypothesis: body size effects on colour-pattern saliency were sex-specific and body size amplifies chromatic saliency in males across different receivers. Large males exhibited high saturation contrast across lizard, bird and snake vision systems (pd _(M>F)_ = 1.00, Figure S4, Table S5), and higher chromatic edge contrast specifically under bird vision (p _(M>F)_ = 0.992). In contrast, females showed non-significant or minimal effects of body size on colour-pattern saliency, with larger females showing higher boundary luminance contrast under lizard and snake vision only, consistent with a cryptic strategy (pd_(M>F)_ < 0.05; Table S4, S5).

### Phylogenetic signal in colour-pattern components is stronger under conspecific and bird vision

Dorsal colour pattern components as perceived by different receivers were phylogenetically conserved across the studied populations, partially supporting our hypothesis (Table S6). Under conspecific and bird vision, most colour-pattern components showed significant phylogenetic signal (−0.51 to 1.06), while snake vision yielded a weaker phylogenetic structure, with only geometric complexity (-0.29) and colour diversity (-0.35) showing significant phylogenetic signal (Table S6). Saturation contrast showed no phylogenetic signal under any visual systems.

## Discussion

Our results suggest that dorsal colour patterns in *P. lusitanicus* resolve the trade-off between conspecific communication and predator detection through receiver- and distance-dependent variation in colour-pattern detectability (Figure 3). Chromatic and achromatic components contribute distinctly to detectability across receivers and viewing distances, with sex-specific receiver strategies and body size amplifying chromatic saliency in males.

**Figure 3.**
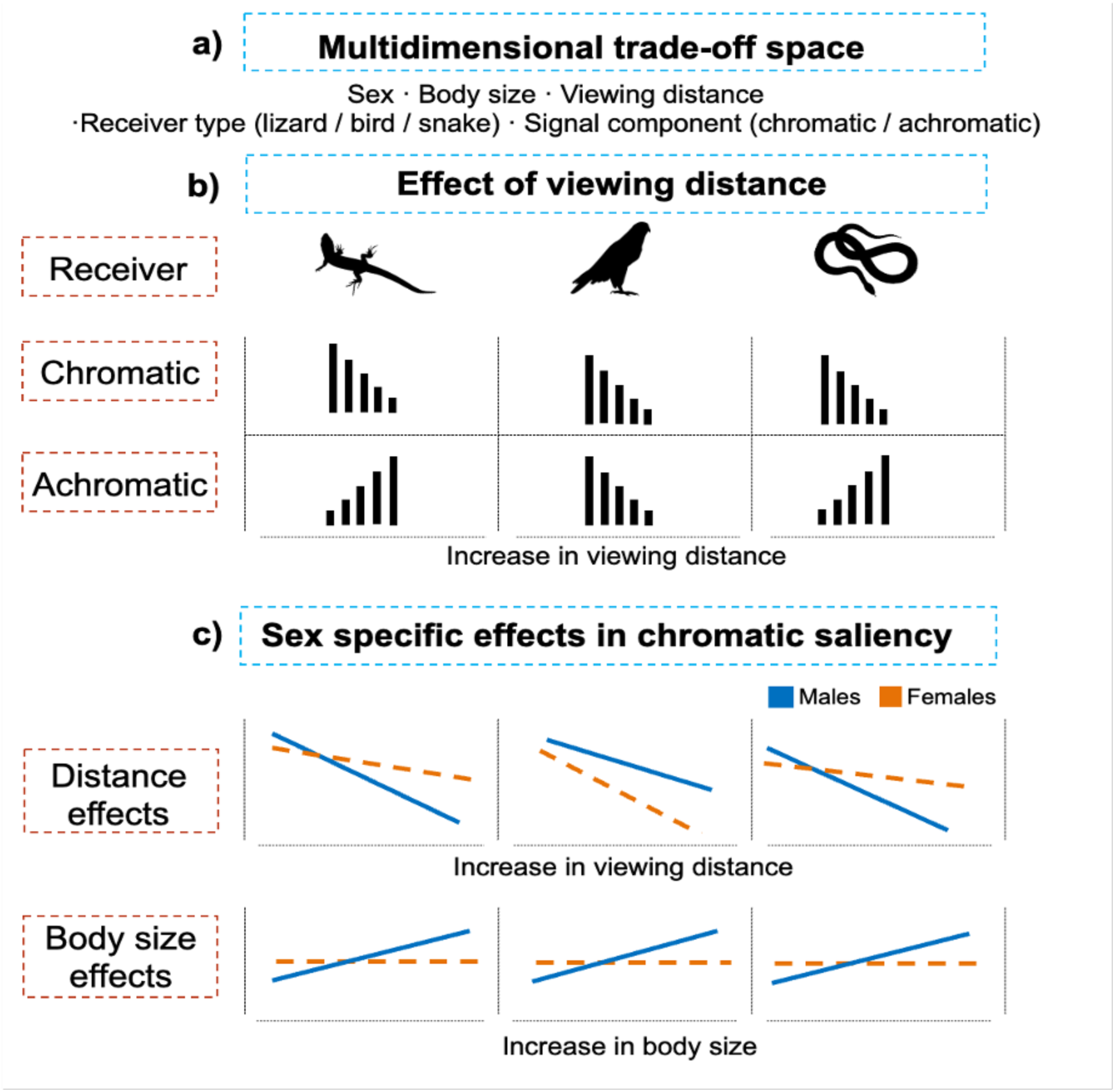
Conceptual summary of how dorsal colour-pattern components of *Podarcis lusitanicus* are perceived across viewing distances by different receiver vision systems. (a) The multidimensional trade-off space defining the key axes of variation shaping colour-pattern perception in this study. (b) Conceptual trends in chromatic and achromatic colour-pattern detectability with increasing viewing distance under conspecific lizard, avian predator, and snake vision systems. The bar plots illustrate the general direction of change in detectability across viewing distances for each receiver. (c) Conceptual summary of sex-specific patterns in chromatic saliency across receivers, showing the effects of viewing distance (top row) and body size (bottom row) in males (blue, solid lines) and females (orange, dashed lines). All silhouettes were obtained from PhyloPic.org. The conspecific lizard silhouette is attributed to Titouan Montessui (licensed under CC BY 4.0, https://creativecommons.org/licenses/by/4.0/).The violet-sensitive bird and snake silhouettes were sourced from openly licensed images.

Colour patterns are predicted to balance conspecific communication against predator avoidance over viewing distance. Our results broadly support this hypothesis but show that the specific trade-offs resolved depend on the receiver’s visual system and viewing distance. Under conspecific lizard and snake vision, chromatic components faded with distance. However, pattern structure and achromatic definition of the boundaries became more prominent (Figures 2 and S3). In the congener *Podarcis muralis*, lizards can detect a conspecific from 10 m (Kawamoto et al., 2025), yet colour patches may blur into the background well within that range (Fleishman & Persons, 2001; Barnett et al., 2018). So, the spatial patterning and luminance boundaries, rather than fine colour details, probably carry the more functionally relevant information for conspecific communication and detection by snake predators at further viewing distances (Katti et al., 2019; Simões et al., 2016), though this channel-resolution difference has yet to be tested directly in lizards. Under bird vision, chromatic and achromatic boundaries became less distinct and fine scale pattern structure decreased at long distance, even though some chromatic metrics (e.g., colour variance and saturation contrast) remained relatively stable (Figures 2 and S3). This may be because the chromatic channel has lower spatial resolution than the achromatic channel (Potier et al., 2018), and low-spatial-frequency signals are generally more robust to blur than fine, high-resolution detail (Mullen, 1985). Consequently, the same dorsal pattern conveys markedly different information to different observers over different viewing distances (Figure 3).

The more information projecting sex, males, resolves trade-offs through distance-dependent colour-pattern detectability. Under conspecific lizard and snake vision, chromatic saliency of male lizards showed greater declines over distance than that of females (Table S5), suggesting that male colour patterns may be optimised for close-distance intraspecific communication, potentially at the cost of increased detectability by snakes. Under bird vision, the pattern is contrasting: detectability of colour variance, boundary luminance contrast, and saturation contrast decline more with distance in females than in males (Table S5). This indicates that both sexes may carry distinct detectability strategies tuned to different observer types, rather than males alone resolving a cost of intraspecific communication (Marshall & Stevens, 2014). This possibility warrants experimental tests quantifying concealment against natural backgrounds (Husak et al., 2006). Although the most visually salient colour patches in wall lizards are typically located on the venter and flanks (Pérez i de Lanuza et al., 2014), sex-specific differences and seasonal changes in dorsal colouration described in several *Podarcis* species (Storniolo et al., 2021) may suggest that males tend to have more salient dorsal colours, possibly contributing to intraspecific communication.

Large individuals lose the resolution of communication-concealment trade-offs by remaining detectable to both conspecifics and predators. Body size has been widely investigated in association with colour-pattern evolution (Goldenberg et al., 2022; Kang et al., 2017; Karpestam et al., 2014). Under conspecific lizard vision, chromatic traits of larger individuals were stronger and more salient at close range (Figure S4). Thus, increasing animal size enhanced chromatic traits for short-range communication while achromatic patterns became the primary information available at greater distances (Bonnaffé et al., 2018; Pérez i de Lanuza & Font, 2015; Ruiz-Miñano et al., 2024). Larger males showed greater overall detectability with higher saturation contrast across all vision systems and higher chromatic edge contrast under bird vision (Table S5, Figure S4; Salisbury & Peters, 2025). Saturation contrast remained relatively stable with distance under bird and snake vision, but declined under lizard vision, indicating that perception of saturation might be more distance-resistant for some receivers in the studied populations. In contrast, larger females exhibited stronger luminance boundaries under lizard and snake vision. Together, these patterns suggest a trade-off in which enhanced chromatic saliency in males may facilitate close-range communication to conspecifics but at the cost of increased visibility to predators in bigger males (Marshall et al., 2015; Marshall & Stevens, 2014; Stuart-Fox et al., 2003). However, studies of congeneric species (*P. muralis* and *P. siculus*) found no sex differences in detectability under kestrel vision, but they did not consider the effect of body size (Costantini & Dell’Omo, 2010). Since larger lizards represent more profitable prey to birds (Costantini et al., 2007; Martin & Lopez, 1990), increased dorsal saliency may primarily be favoured in large, dominant males when communication benefits outweigh predation costs. The fitness costs of this trade-off will require experimental tests of predation risk using prey models (Amdekar & Thaker, 2019; Husak et al., 2006). Finally, body size and sex effects in colour-pattern detectability also varied among populations (Table S3), indicating that local ecological conditions may further shape colour pattern variation (Stuart-Fox & Ord, 2004).

The strength of phylogenetic signals in colour-pattern traits is the strongest under conspecific and bird vision (5/6 and 4/6 traits, respectively) and weakest under snake vision (2/6 traits; Table S6). Traits relevant for conspecific communication and avian predation may be evolutionarily conserved through stabilising selection, reflecting the broad and consistent selective pressures imposed by intraspecific interactions and dominant in NW Iberia predators of wall lizards (Allen et al., 2020; Martin & Lopez, 1990; Pérez i de Lanuza & Font, 2016). Snakes, as ectotherms, have substantially lower energetic demands than endothermic birds, may consume less prey biomass per individual, and thus representing much less selective force on prey population (Nagy, Girard & Brown, 1999) and may therefore impose weaker predator-mediated selection on prey visual traits. Also, the use of chemoreception by snakes for accurate prey detection may reduce the selective importance of prey dorsal colouration (Revell et al., 2008; Saviola et al., 2012). This is consistent with our results of lizards tolerating increased visibility to snakes at short distances when it enhances conspecific communication. Saturation contrast, that showed no phylogenetic signal under all vision systems, may be evolutionary unconstrained reflecting potential high evolutionary responsiveness to variable local environmental factors, such as predation, micro-habitat conditions or thermoregulatory demands (Freitas et al., 2021; Murali et al., 2024; Koneru & Caro, 2022). Nevertheless, our results suggest the importance of both communication and predation shaping dorsal colour patterns in wall lizards.

While our results indicate that dorsal colour patterns comprise multiple components whose visual effectiveness vary across spatial scales and receiver types, (Harper, 2006; Nokelainen et al., 2022; Pérez-Rodríguez et al., 2017; Macedo et al., 2022), they should be interpreted with caution, as the colour-pattern outputs derive from the same reflectance data, but are processed through vision models that differ in photoreceptor dimensionality and noise structure. Despite extensive sampling across populations and individuals, our phylogenetic analysis was limited by the restricted geographic range and shallow phylogeographic structure of *P. lusitanicus* (Rato et al., 2024), potentially constraining our ability to detect deeper patterns of trait covariation that may emerge in more species with broader ecological and geographic ranges with deeper phylogenetic sub structuring. We recommend behavioural assays of prey detection and conspecific responses under ecologically realistic light and background conditions, particularly for high-acuity avian predators, to test whether the achromatic saliency patterns observed under bird vision persist over greater viewing distances. Finally, survival experiments are needed to determine whether the perceptual changes we measured translate into differences in detection, discrimination and fitness (Marshall et al., 2015, 2016; Purger et al., 2017; Ruiz-Miñano et al., 2024).

## Conclusions

Our study reveals that the detectability of dorsal colour-pattern components in Lusitanian wall lizards is not tuned to a single optimum but instead reflects a compromise among multiple, often contrasting forces. The interplay between viewing distance, observer visual system and body size reflects sex-specific trade-offs between conspecific communication and detectability by predators. Detectability shifted from chromatic information to achromatic pattern structure with increasing distance under conspecific and snake vision, while both components declined uniformly under bird vision, revealing fundamentally different perceptual landscapes across observer types. Sex- and size-specificity of colour-pattern saliency suggest that this trade-off is resolved differently across individuals rather than converging on a common optimum. In females and small individuals, colour patterns appear constrained towards reduced saliency, consistent with a strategy that prioritises predator avoidance when the benefits of salient colouration are limited. This pattern is lost in large males that are salient despite high risk of predation. As our data address perceptual salience rather than realised survival or mating outcomes, we cannot directly assess the fitness consequences of detected patterns. Nonetheless, these findings support the view that colour signals evolve under multi-receiver selection, with the balance between intraspecific communication and predator avoidance. These findings underscore the value of multi-receiver, distance-dependent frameworks for understanding how conflicting selective pressures jointly shape colour-pattern evolution.

## Ethics Statement

All applicable international, national and/or institutional guidelines for the care and use of animals were followed. Collecting permits 536 / 2022 / CAPT and EB-042/2023 were provided by the Institute for Nature Conservation and Forests (ICNF, Portugal) and the Xunta de Galicia (Spain), respectively. This study was approved by the ethical guidelines of the University of Porto.

## Data accessibility

The data and code supporting the findings of this study will be made public upon acceptance of the manuscript

## Acknowledgements

We thank Dr. C. van den Berg for valuable guidance on the Batch QCPA analyses during the development of this work. We are grateful to L. Papaleo, I. Dannecker, L. Gautier, I. Ferreira, G. Simbula, L. Santos, G. Fănaru, G. Ene, A. Limnios, J. Faria, and C. Faria for their assistance with fieldwork and photographic sampling. We also thank César Ayres for providing detailed locality information for the Spanish populations. This work was supported by Fundação para a Ciência e Tecnologia (FCT, Portugal) through the projects PTDC/BIA-CBI/28014/2017 and 2022.03391.PTDC. LBS was supported by a PhD grant (2021.06600.BD) from FCT.

