## Supplementary file for "Resolution of multi-receiver trade-offs in wall lizard colouration"

**Supplementary material**

**Table S1.** Visual system parameters used for visual system modelling. We set the acuity of conspecific lizard model to 2.05 cpd (*Podarcis muralis**, Kawamoto et al., 2025), bird predator to 40cpd (*Falco sparverius**, Hirsch, 1982) and snake predator to 4 cpd (*Nerodia sipedon pleuralis*,* Baker et al., 2007). LWS: long-wavelength-sensitive; MWS: medium-wavelength-sensitive; SWS: short-wavelength sensitive; VS: violet- sensitive; UVS: ultraviolet-sensitive. Weber fraction estimates of 0.05 in the absence of specific data as for snake vision, (Salisbury & Peters, 2025; Vorobyev et al., 2001).

| Vision models used | Cone type | λmax | Weber fraction | Citation |
|---|---|---|---|---|
| Wall lizard | *LWS* | 562 | 0.05 | Martin et al., 2015 |
| (*Podarcis muralis*) | *MWS* | 497 | 0.1 | |
| (Cone ratio: 1:1:1:4) | *SWS* | 456 | 0.1 | |
| | *UVS* | 367 | 0.1 | |
| Peafowl | *LWS* | 599 | 0.051 | Hart., 2002 |
| (*Pavo cristatus*) | *MWS* | 537 | 0.05 | |
| (Cone ratio: 1 : 1.9 : 2.2 : 2.1) | *SWS* | 477 | 0.054 | |
| | *VS* | 433 | 0.074 | |
| | *double cone* | 567 | 0.1 | |
| Garter snake | *LWS* | 554 | 0.1 | Goedert et al., 2021; Sillman et al., 1997 |
| (*Thamnophis sirtalis*) | *MWS* | 482 | 0.05 | |
| (Cone ratio: 1:1.6:7.3) | *SWS1* | 360 | 0.05 | |

*In the absence of data for the conspecific/predator species of *Podarcis lusitanicus*, we used values of other species that were phylogenetically close.

**Table S2**. Summary statistics of the colour-pattern variables of the dorsum of *Podarcis lusitanicus*, modelled for three different vision systems: 1) conspecific lizard, 2) bird predator 3) snake predator

| Modelled vision system | Sex | Variable | Mean | SD | Skewness | Kurtosis |
|---|---|---|---|---|---|---|
| Lizard vision | Male (N= 262) | Chromatic edge contrast | 0.05 | 0.98 | 1.42 | 4.49 |
| | | Colour variance | -0.02 | 1.05 | -0.51 | 2.9 |
| | | Geometric complexity | 0.04 | 1.01 | 0.68 | 3.47 |
| | | Colour diversity | 0.07 | 1.02 | 0.09 | 2.94 |
| | | Saturation contrast | 0.16 | 0.93 | -0.34 | 4.13 |
| | | Boundary luminance contrast | -0.07 | 0.96 | 0.36 | 2.33 |
| | Female (N= 201) | Chromatic edge contrast | -0.1 | 0.86 | 1.68 | 5.73 |
| | | Colour variance \| | 0.02 | 0.93 | -0.41 | 2.77 |
| | | Geometric complexity | -0.05 | 0.99 | 0.67 | 3.58 |
| | | Colour diversity | -0.09 | 0.96 | 0.09 | 2.74 |
| | | Saturation contrast | -0.25 | 0.94 | -0.45 | 4.19 |
| | | Boundary luminance contrast | 0.08 | 1.04 | 0.42 | 2.54 |
| Bird vision | Male (N= 261) | Chromatic edge contrast | 0.01 | 1.01 | 1.01 | 3.56 |
| | | Colour variance | -0.07 | 1.05 | 0.62 | 4.04 |
| | | Geometric complexity | 0.04 | 1.03 | 0.26 | 2.4 |
| | | Colour diversity | 0.06 | 1.01 | 0.44 | 2.88 |
| | | Saturation contrast | 0.15 | 0.87 | -0.13 | 3.53 |

| | | | | | | |
|---|---|---|---|---|---|---|
| | | Boundary luminance contrast | -0.06 | 0.94 | -0.07 | 3.94 |
| | Female (N= 202) | Chromatic edge contrast | -0.02 | 0.99 | 0.85 | 3.14 |
| | | Colour variance | 0.09 | 0.91 | 0.48 | 3.57 |
| | | Geometric complexity | -0.04 | 0.96 | 0.03 | 2.18 |
| | | Colour diversity | -0.07 | 0.98 | 0.49 | 2.73 |
| | | Saturation contrast | -0.24 | 0.91 | 0.15 | 4.92 |
| | | Boundary luminance contrast | 0.09 | 1.07 | -0.16 | 3.14 |
| Snake vision | Male (N= 261) | Chromatic edge contrast | 0.03 | 0.9 | 1.33 | 5.09 |
| | | Colour variance \| | 0.01 | 1.04 | -0.49 | 3.16 |
| | | Geometric complexity | 0.02 | 1.03 | 1.13 | 4.8 |
| | | Colour diversity | 0.02 | 1 | 0.78 | 3.22 |
| | | Saturation contrast | 0.16 | 0.9 | 0.07 | 3.61 |
| | | Boundary luminance contrast | -0.12 | 0.95 | 0.13 | 2.78 |
| | Female (N= 200) | Chromatic edge contrast | -0.12 | 0.79 | 1.4 | 5.44 |
| | | Colour variance | -0.02 | 0.93 | -0.38 | 3.34 |
| | | Geometric complexity | -0.02 | 0.97 | 0.88 | 3.85 |
| | | Colour diversity | -0.05 | 0.95 | 0.81 | 3.06 |
| | | Saturation contrast | -0.16 | 0.94 | -0.11 | 3.42 |
| | | Boundary luminance contrast | 0.16 | 1.03 | 0.6 | 4.34 |

**Table S3.** Results of Multivariate analysis of variance (MANOVA) testing the effects of population, sex, body size, and their interactions on dorsal colour-pattern components of *Podarcis lusitanicus* under conspecific (lizard) and predator (snake, violet-sensitive bird) vision systems for a viewing distance of 1.5m. For each effect, Pillai's trace, F-statistics with numerator and denominator degrees of freedom, and associated p-values are reported.

| Vision system | Effect | Df | Pillai | F $_{(num\ df,\ den\ df)}$ | P |
|---|---|---|---|---|---|
| Lizard (Conspecific) | Population | 20 | 1.3 | 5.19 $_{(120,\ 2262)}$ | <0.001 |
| | Sex | 1 | 0.18 | 13.88 $_{(6,\ 372)}$ | <0.001 |
| | Body size | 1 | 0.31 | 28.14 $_{(6,\ 372)}$ | <0.001 |
| | Population: Sex | 20 | 0.38 | 1.26 $_{(120,\ 2262)}$ | <0.05 |
| | Population: Body size | 20 | 0.39 | 1.30 $_{(120,\ 2262)}$ | <0.05 |
| | Sex: Body size | 1 | 0.07 | 4.58 $_{(6,\ 372)}$ | <0.001 |
| Bird (predator) | Population | 20 | 2 | 9.73 $_{(120,\ 2334)}$ | <0.001 |
| | Sex | 1 | 0.24 | 20.52 $_{(6,\ 384)}$ | <0.001 |
| | Body size | 1 | 0.2 | 16.13 $_{(6,\ 384)}$ | <0.001 |
| | Population: Sex | 20 | 0.42 | 1.46 $_{(120,\ 2334)}$ | <0.01 |
| | Population: Body size | 20 | 0.39 | 1.35 $_{(120,\ 2334)}$ | <0.01 |
| | Sex: Body size | 1 | 0.06 | 3.98 $_{(6,\ 384)}$ | <0.01 |
| Snake (predator) | Population | 20 | 1.15 | 4.62 $_{(120,\ 2334)}$ | <0.001 |
| | Sex | 1 | 0.36 | 35.45 $_{(6,\ 384)}$ | <0.001 |
| | Body size | 1 | 0.4 | 42.22 $_{(6,\ 384)}$ | <0.001 |
| | Population: Sex | 20 | 0.36 | 1.25 $_{(120,\ 2334)}$ | <0.05 |
| | Sex: Body size | 1 | 0.06 | 3.95 $_{(6,\ 384)}$ | <0.01 |

**Table S4.** Summary of the effects of viewing distance and body length on the colour-pattern metrics of the dorsal appearance of *Podarcis lusitanicus* lizards across different observer vision systems. For each vision system, colour-pattern metrics derived from the quantitative colour-pattern analysis (QCPA) were modelled using Bayesian multivariate mixed models, with viewing distance and body size as predictors (15,000 iterations, 1,000 warm-up, thinning = 10, with weakly informative priors). Males and females were analysed separately. Individual lizards nested in sampling location is added as random factor. Estimates are reported as mean ± 95% credible intervals (CI). Viewing distances modelled: 1) conspecific lizard: 0.5m (short), 1.5m (medium), 2m (long); 2) Bird predator: 1.5m (short), 3m (medium), 4m (long); (3) snake predator: 1m (short), 1.5m (medium), 3m (long).

| Response variable | Predictor | Lizard vision (Estimate ± 95% CI) | | Bird vision (Estimate ± 95% CI) | | Snake vision (Estimate ± 95% CI) | |
|---|---|---|---|---|---|---|---|
| | | Male | Female | Male | Female | Male | Female |
| Chromatic edge contrast (Col.mean) | Intercept | 1.14 (1.05, 1.23) | 0.87 (0.78, 0.96) | 1.11 (1.01, 1.22) | 1.09 (0.97, 1.20) | 0.84 (0.75, 0.94) | 0.63 (0.54, 0.72) |
| | Distance (medium) | −1.66 (−1.74, −1.58) | −1.42 (−1.52, −1.33) | –1.57 (–1.65, –1.50) | –1.51 (–1.59, –1.44) | –0.73 (–0.83, –0.64) | –0.69 (–0.78, –0.60) |
| | Distance (long) | −1.87 (−1.96, −1.79) | −1.57 (−1.67, −1.47) | –1.90 (–1.98, –1.83) | –1.89 (–1.97, –1.81) | –1.62 (–1.72, –1.53) | –1.43 (–1.52, –1.33) |
| | Body size | 0.17 (0.13, 0.21) | 0.14 (0.09, 0.18) | 0.21 (0.17, 0.25) | 0.12 (0.06, 0.18) | 0.15 (0.10, 0.21) | 0.13 (0.06, 0.19) |

| | | | | | | | |
|---|---|---|---|---|---|---|---|
| Colour variance (Col.CoV) | Intercept | 0.79<br>(0.57, 1.02) | 0.69<br>(0.52, 0.87) | –0.43<br>(–0.73, –0.15) | –0.01<br>(–0.21, 0.19) | 0.79<br>(0.59, 1.00) | 0.71<br>(0.54, 0.88) |
| | Distance (medium) | −1.20<br>(−1.29, −1.12) | −1.00<br>(−1.09, −0.90) | 0.62<br>(0.53, 0.70) | 0.29<br>(0.18, 0.40) | –0.63<br>(–0.75, –0.51) | –0.69<br>(–0.80, –0.59) |
| | Distance (long) | −1.24<br>(−1.34, −1.15) | −0.96<br>(−1.06, -0.85) | 0.55<br>(0.47, 0.63) | 0.15<br>(0.04, 0.27) | –1.51<br>(–1.63, –1.40) | –1.30<br>(–1.41, –1.19) |
| | Body size | 0.12<br>(0.04, 0.20) | 0.10<br>(0.00, 0.20) | 0.10<br>(0.00, 0.21) | 0.08<br>(–0.04, 0.20) | 0.22<br>(0.14, 0.30) | 0.19<br>(0.09, 0.29) |
| Geometric complexity (CAA.C) | Intercept | −0.40<br>(−0.54, −0.27) | −0.38<br>(−0.52, −0.25) | 0.93<br>(0.78, 1.06) | 0.72<br>(0.54, 0.89) | –0.41<br>(–0.55, –0.28) | –0.67<br>(–0.82, –0.52) |
| | Distance (medium) | 0.87<br>(0.72, 1.02) | 0.61<br>(0.43, 0.79) | –1.37<br>(–1.49, –1.24) | –1.14<br>(–1.27, –1.01) | –0.02<br>(–0.18, 0.13) | 0.55<br>(0.38, 0.71) |
| | Distance (long) | 0.42<br>(0.25, 0.58) | 0.38<br>(0.18, 0.58) | –1.51<br>(–1.63, –1.39) | –1.29<br>(–1.42, –1.15) | 1.24<br>(1.08, 1.40) | 1.33<br>(1.16, 1.50) |
| | Body size | 0.06<br>(−0.02, 0.13) | −0.03<br>(−0.11, 0.06) | 0.14<br>(0.08, 0.19) | 0.10<br>(0.03, 0.16) | –0.02<br>(–0.09, 0.04) | –0.06<br>(–0.14, 0.01) |
| Colour diversity | Intercept | −0.40<br>(−0.55, −0.26) | −0.44<br>(−0.60, −0.29) | 0.69<br>(0.49, 0.89) | 0.63<br>(0.41, 0.85) | –0.05<br>(–0.20, 0.10) | –0.41<br>(–0.56, –0.24) |

| | | | | | | | |
|---|---|---|---|---|---|---|---|
| (CAA.Sc) | Distance (medium) | 0.91 (0.76, 1.06) | 0.79 (0.62, 0.96) | –0.99 (–1.10, –0.88) | –1.08 (–1.20, –0.97) | –0.49 (–0.66, –0.32) | 0.21 (0.04, 0.38) |
| | Distance (long) | 0.30 (0.14, 0.46) | 0.25 (0.07, 0.42) | –1.23 (–1.34, –1.13) | –1.24 (–1.36, –1.12) | 0.56 (0.39, 0.72) | 0.83 (0.65, 1.01) |
| | Body size | 0.21 (0.14, 0.28) | 0.15 (0.07, 0.23) | 0.14 (0.07, 0.20) | 0.13 (0.06, 0.19) | 0.09 (0.03, 0.16) | 0.07 (–0.02, 0.15) |
| Saturation contrast (VCA.MSsat) | Intercept | 0.21 (−0.10, 0.51) | −0.19 (−0.52, 0.12) | 0.07 (–0.20, 0.32) | –0.25 (–0.52, 0.03) | 0.04 (–0.26, 0.34) | –0.33 (–0.66, 0.01) |
| | Distance (medium) | −0.19 (−0.23, −0.16) | −0.12 (−0.16, −0.08) | 0.03 (0.01, 0.05) | 0.01 (0.00, 0.03) | 0.02 (–0.01, 0.05) | 0.05 (0.02, 0.09) |
| | Distance (long) | −0.26 (−0.30, −0.23) | −0.17 (−0.22, −0.13) | 0.05 (0.03, 0.06) | 0.02 (0.01, 0.04) | 0.15 (0.12, 0.19) | 0.26 (0.23, 0.29) |
| | Body size | 0.21 (0.12, 0.29) | −0.09 (−0.20, 0.01) | 0.21 (0.12, 0.30) | 0.01 (–0.12, 0.14) | 0.15 (0.06, 0.23) | –0.12 (–0.21, –0.03) |
| Boundary luminance contrast (BSA.BML) | Intercept | −0.95 (−1.06, −0.85) | −0.84 (−0.95, −0.73) | 0.27 (0.01, 0.53) | 0.58 (0.32, 0.84) | –0.96 (–1.08, –0.83) | –0.56 (–0.71, –0.41) |
| | Distance (medium) | 1.28 (1.16, 1.39) | 1.48 (1.33, 1.63) | –0.49 (–0.61, –0.38) | –0.67 (–0.81, –0.53) | 0.82 (0.69, 0.96) | 0.69 (0.51, 0.88) |

| | | | | | | |
|---|---|---|---|---|---|---|
| Distance (long) | 1.50 (1.38, 1.62) | 1.42 (1.25, 1.57) | –0.66 (–0.78, –0.55) | –0.96 (–1.11, –0.81) | 1.55 (1.42, 1.69) | 1.40 (1.21, 1.59) |
| Body size | −0.07 (−0.13, −0.02) | 0.02 (−0.05, 0.08) | 0.09 (0.02, 0.16) | 0.09 (–0.01, 0.19) | –0.17 (–0.23, –0.11) | –0.07 (–0.15, 0.01) |

Body size effects were sex- and vision system-dependent, enhancing multiple colour pattern traits in males under both conspecific and predator vision systems, while in females, effects were weaker under lizard vision and reduced or absent under predator vision (Figure 2, Figure S4, Table S4). Positive effects emerged for males in chromatic edge contrast ($\beta$ = lizard 0.17, snake 0.15), colour variance (lizard 0.12, snake 0.22), colour diversity (lizard 0.21, snake 0.09), and saturation contrast (lizard 0.21, snake 0.15); though boundary luminance contrast decreased in snake vision (−0.17). In females, body size effects were more limited, with positive effects on chromatic edge contrast (lizard 0.14, snake 0.13), colour diversity (lizard 0.15) and colour variance (snake 0.19). Under snake vision, saturation contrast decreased (−0.12) with body size in females, contrasting with the positive male effect (Figure S4). In bird vision, body size had positive effects on most colour pattern descriptors in both sexes (Figure S4); the strongest effects observed for chromatic edge contrast (males 0.21, females 0.12) and saturation contrast (only males 0.21).

**Table S5.** Sex-specific differences in the effects of body length and viewing distance on colour-pattern variables in Lusitanian wall lizard (*Podarcis lusitanicus*) under different observer vision systems; lizard (conspecific), bird (predator) and snake (predator). For each response variable and predictor, the table reports the percentage of overlap of the posterior distributions between sexes; both total (Overlap total %) and within 95% credible intervals (Overlap CI %). Median posterior differences between male and female effect sizes are reported with 95% highest density intervals (HDIs); differences are considered credible when the 95% HDI does not include zero. Probability of direction (pd) with the direction of the effect (F>M or M>F) is reported; values approaching 1.0 indicates high certainty that males show stronger effects, while values approaching 0.0 indicate females show stronger effects.

Viewing distances modelled: 1) conspecific lizard: 0.5m (short), 1.5m (medium), 2m (long); 2) Bird predator: 1.5m (short), 3m (medium), 4m (long); (3) snake predator: 1m (short), 1.5m (medium), 3m (long).

| Vision system | Response variable | Predictor | Overlap total % (Overlap CI %) | Overlap interpretation* | Median difference (HDI lower, HDI upper) | Difference status | pd (direction) |
|---|---|---|---|---|---|---|---|
| Lizard (conspecific) | Boundary luminance contrast | Body length | 44.13 (25.01) | Minimal | -0.09 (-0.18, 0.00) | Credible | 0.024 (F > M) |
| | | Distance: Medium vs Short | 43.84 (22.75) | Minimal | -0.20 (-0.39, -0.01) | Credible | 0.019 (F > M) |

| | | | | | | |
|---|---|---|---|---|---|---|
| | Distance: Long vs Short | 79.55 (68.49) | Substantial | 0.09 (-0.12, 0.29) | Not credible | 0.804 (M > F) |
| Chromatic edge contrast | Body length | 65.38 (60.13) | Substantial | 0.03 (-0.03, 0.09) | Not credible | 0.864 (M > F) |
| | Distance: Medium vs Short | 18.55 (0.00) | No | -0.24 (-0.36, -0.11) | Credible | 0.001 (F > M) |
| | Distance: Long vs Short | 22.83 (0.00) | No | -0.30 (-0.43, -0.17) | Credible | 0.000 (F > M) |
| Colour diversity | Body length | 66.46 (58.99) | Substantial | 0.06 (-0.05, 0.17) | Not credible | 0.864 (M > F) |
| | Distance: Medium vs Short | 79.74 (63.16) | Substantial | 0.12 (-0.11, 0.34) | Not credible | 0.849 (M > F) |
| | Distance: Long vs Short | 81.01 (84.38) | High | 0.05 (-0.19, 0.29) | Not credible | 0.647 (M > F) |
| Colour variance | Body length | 79.40 (87.41) | High | 0.03 (-0.10, 0.15) | Not credible | 0.650 (M > F) |
| | Distance: Medium vs Short | 20.75 (0.00) | No | -0.21 (-0.34, -0.08) | Credible | 0.001 (F > M) |
| | Distance: Long vs Short | 13.22 (0.00) | No | -0.29 (-0.43, -0.15) | Credible | 0.000 (F > M) |

| | | | | | | | |
|---|---|---|---|---|---|---|---|
| | Geometric complexity | Body length | 61.43 (47.18) | Moderate | 0.08 (-0.03, 0.19) | Not credible | 0.931 (M > F) |
| | | Distance: Medium vs Short | 44.46 (23.03) | Minimal | 0.26 (0.01, 0.50) | Credible | 0.981 (M > F) |
| | | Distance: Long vs Short | 84.62 (90.11) | High | 0.03 (-0.22, 0.30) | Not credible | 0.594 (M > F) |
| | Saturation contrasts | Body length | 11.36 (0.00) | No | 0.30 (0.16, 0.44) | Credible | 1.000 (M > F) |
| | | Distance: Medium vs Short | 31.63 (7.64) | Minimal | -0.07 (-0.13, -0.02) | Credible | 0.005 (F > M) |
| | | Distance: Long vs Short | 27.48 (0.00) | No | -0.09 (-0.15, -0.03) | Credible | 0.002 (F > M) |
| Bird (predator) | Boundary luminance contrast | Body length | 70.55 (81.59) | High | 0.01 (-0.13, 0.12) | Not credible | 0.494 (F > M) |
| | | Distance: Medium vs Short | 53.23 (28.94) | Moderate | 0.18 (0.00, 0.36) | Not credible | 0.974 (M > F) |
| | | Distance: Long vs Short | 22.52 (0.00) | No | 0.30 (0.11, 0.49) | Credible | 0.999 (M > F) |
| | Chromatic edge contrast | Body length | 31.59 (9.87) | Minimal | 0.09 (0.02, 0.17) | Credible | 0.992 (M > F) |

| | | | | | | |
|---|---|---|---|---|---|---|
| | Distance: Medium vs Short | 65.81 (59.59) | Substantial | -0.06 (-0.17, 0.05) | Not credible | 0.151 (F > M) |
| | Distance: Long vs Short | 94.78 (89.82) | High | -0.02 (-0.12, 0.09) | Not credible | 0.392 (F > M) |
| Colour diversity | Body length | 87.27 (92.69) | High | 0.01 (-0.08, 0.10) | Not credible | 0.581 (M > F) |
| | Distance: Medium vs Short | 58.43 (57.63) | Substantial | 0.10 (-0.06, 0.25) | Not credible | 0.876 (M > F) |
| | Distance: Long vs Short | 90.11 (92.19) | High | 0.01 (-0.15, 0.18) | Not credible | 0.555 (M > F) |
| Colour variance | Body length | 95.27 (89.03) | High | 0.02 (-0.13, 0.18) | Not credible | 0.607 (M > F) |
| | Distance: Medium vs Short | 8.95 (0.00) | No | 0.33 (0.19, 0.46) | Credible | 1.000 (M > F) |
| | Distance: Long vs Short | 0.00 (0.00) | No | 0.40 (0.26, 0.54) | Credible | 1.000 (M > F) |
| Geometric complexity | Body length | 77.31 (68.41) | Substantial | 0.04 (-0.05, 0.12) | Not credible | 0.807 (M > F) |
| | Distance: Medium vs Short | 36.01 (11.36) | Minimal | -0.23 (-0.41, -0.05) | Credible | 0.005 (F > M) |

| | | | | | | | |
|---|---|---|---|---|---|---|---|
| | | Distance: Long vs Short | 38.08 (13.86) | Minimal | -0.22 (-0.41, -0.04) | Credible | 0.009 (F > M) |
| | Saturation contrasts | Body length | 36.92 (9.43) | Minimal | 0.20 (0.04, 0.36) | Credible | 0.993 (M > F) |
| | | Distance: Medium vs Short | 48.96 (42.32) | Moderate | 0.02 (0.00, 0.04) | Not credible | 0.941 (M > F) |
| | | Distance: Long vs Short | 38.42 (14.90) | Minimal | 0.02 (0.00, 0.04) | Credible | 0.990 (M > F) |
| Snake (Predator) | Boundary luminance contrast | Body length | 43.11 (25.92) | Moderate | -0.10 (-0.20, 0.00) | Credible | 0.021 (F > M) |
| | | Distance: Medium vs Short | 60.37 (59.80) | Substantial | 0.13 (-0.09, 0.35) | Not credible | 0.868 (M > F) |
| | | Distance: Long vs Short | 60.95 (52.78) | Substantial | 0.16 (-0.08, 0.39) | Not credible | 0.904 (M > F) |
| | Chromatic edge contrast | Body length | 75.97 (79.37) | High | 0.03 (-0.06, 0.11) | Not credible | 0.717 (M > F) |
| | | Distance: Medium vs Short | 70.46 (75.27) | High | -0.04 (-0.17, 0.08) | Not credible | 0.256 (F > M) |
| | | Distance: Long vs Short | 31.61 (0.00) | No | -0.20 (-0.33, -0.07) | Credible | 0.002 (F > M) |

| | | | | | | |
|---|---|---|---|---|---|---|
| Colour diversity | Body length | 75.02 (78.28) | High | 0.03 (-0.08, 0.14) | Not credible | 0.708 (M > F) |
| | Distance: Medium vs Short | 0.00 (0.00) | No | -0.71 (-0.94, -0.46) | Credible | 0.000 (F > M) |
| | Distance: Long vs Short | 38.68 (20.01) | Minimal | -0.28 (-0.52, -0.03) | Credible | 0.015 (F > M) |
| Colour variance | Body length | 87.84 (85.36) | High | 0.03 (-0.10, 0.15) | Not credible | 0.664 (M > F) |
| | Distance: Medium vs Short | 64.53 (72.39) | Substantial | 0.06 (-0.10, 0.22) | Not credible | 0.769 (M > F) |
| | Distance: Long vs Short | 36.54 (7.24) | Minimal | -0.21 (-0.38, -0.05) | Credible | 0.007 (F > M) |
| Geometric complexity | Body length | 75.53 (71.68) | Substantial | 0.04 (-0.06, 0.14) | Not credible | 0.786 (M > F) |
| | Distance: Medium vs Short | 8.08 (0.00) | No | -0.57 (-0.79, -0.35) | Credible | 0.000 (F > M) |
| | Distance: Long vs Short | 71.74 (73.07) | Substantial | -0.09 (-0.33, 0.14) | Not credible | 0.219 (F > M) |
| Saturation contrasts | Body length | 11.76 (0.00) | No | 0.27 (0.15, 0.39) | Credible | 1.000 (M > F) |

| | | | | | |
|---|---|---|---|---|---|
| | Distance: Medium vs Short | 48.74 (48.74) | Moderate | -0.03 (-0.08, 0.01) | Not credible | 0.075 (F > M) |
| | Distance: Long vs Short | 0.00 (0.00) | No | -0.11 (-0.15, -0.06) | Credible | 0.000 (F > M) |

*Interpretation of the percentage overlap of the posterior distributions of males and females: 0% = no overlap, <25% = minimal overlap, 25–50% = moderate overlap, 50–75% = substantial overlap, and >75% = high overlap.

Overlap ranged from 13.22–44.46% under lizard vision, 0–25.92% under snake vision, and 0–38.42% under bird vision - varying with viewing distance and body size of lizards (Table S5). Credible sex differences in body size effects on colour pattern components emerged for specific traits, showing minimal posterior overlap between sexes: boundary luminance contrast (posterior overlap%: lizard 44.13%, snake 25.92%), saturation contrast (lizard 11.36%, bird 36.92%, snake 0%), and chromatic edge contrast (bird 31.59%, Table S5).

**Table S6.** Posterior means, confidence intervals, and p-value for regressions testing the phylogenetic signal in the colour-pattern components of Podarcis lusitanicus wall lizards. Population level averages (N= 21) of the colour-pattern components modelled for a distance of 1.5m, across conspecific (lizards) and predators (snakes, violet-sensitive bird) were used as response variables in generalised linear mixed models (glmm) using Markov chain Monte Carlo (200,000 iterations, 8,000 burn-in, thinning = 100) with an intercept only fixed effect structure and population genetic distances as random factor.

| Vision system | Response | Mean | pMCMC (CI lower, CI upper) | ESS | DIC |
|---|---|---|---|---|---|
| Lizard (Conspecific) | Chromatic edge contrast | -0.51 | <0.01 (-0.61, -0.39) | 1920 | -23.3 |
| | Colour variance | -0.36 | <0.05 (-0.71, -0.06) | 2189 | 27.5 |
| | Geometric complexity | 0.33 | <0.05 (0.05, 0.55) | 2299 | 3.8 |
| | Colour diversity | 0.42 | <0.01 (0.26, 0.59) | 1920 | -12.7 |
| | Saturation contrast | -0.12 | 0.620 (-0.70, 0.31) | 1920 | 48.3 |
| | Boundary luminance contrast | 0.49 | <0.01 (0.25, 0.74) | 1920 | 2.45 |
| Bird (Predator) | Chromatic edge contrast | 1.06 | <0.01 (0.70, 1.38) | 1920 | 27.3 |
| | Colour variance | -0.26 | 0.075 (-0.60, 0.02) | 1920 | 18.9 |

| | | | | | |
|---|---|---|---|---|---|
| | Geometric complexity | 0.83 | <0.01 (0.50, 1.16) | 1920 | 25.7 |
| | Colour diversity | 0.66 | <0.01 (0.35, 0.99) | 1920 | 31.7 |
| | Saturation contrast | -0.05 | 0.860 (-0.69, 0.63) | 1650 | 38.5 |
| | Boundary luminance contrast | 0.39 | <0.05 (0.04, 0.70) | 1920 | 20.9 |
| Snake (Predator) | Chromatic edge contrast | 0.1 | 0.272 (-0.12, 0.34) | 1920 | -2.9 |
| | Colour variance | 0.1 | 0.298 (-0.14, 0.34) | 1920 | 3.86 |
| | Geometric complexity | -0.29 | <0.05 (-0.57, -0.03) | 1920 | 10.5 |
| | Colour diversity | -0.35 | <0.05 (-0.61, -0.07) | 1920 | 15.0 |
| | Saturation contrast | -0.12 | 0.579 (-0.58, 0.43) | 1920 | 49.4 |
| | Boundary luminance contrast | -0.01 | 0.912 (-0.22, 0.21) | 1920 | 7.96 |

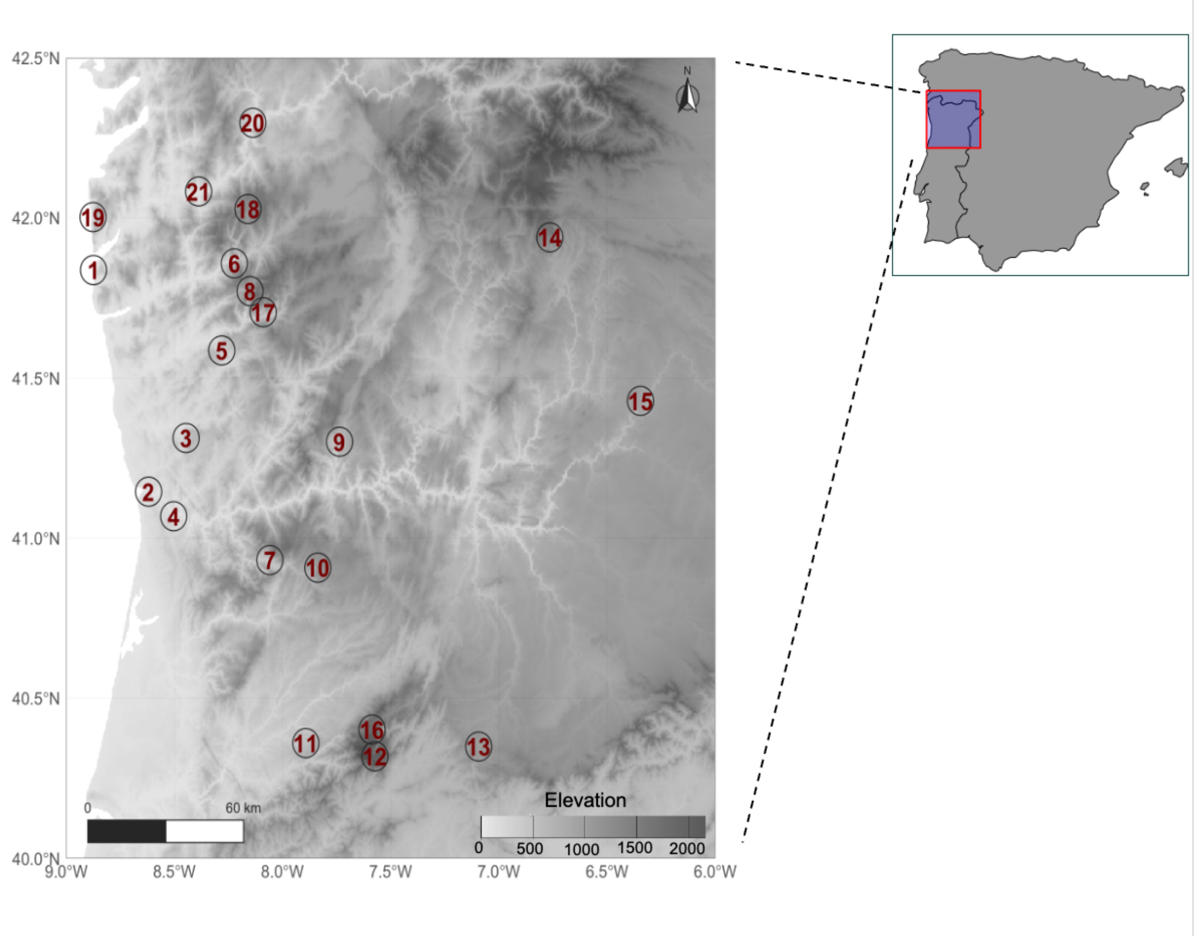


**Figure S1.** Sampling locations and multivariate colour-pattern variation in *Podarcis lusitanicus* across receiver vision systems. Map of the 21 sampling populations across north-western Iberia, overlaid on a digital elevation model (SRTM 30 m resampled to 300 m resolution). The populations sampled and their coordinates (latitude, longitude): 1. Moledo (41.837°N, 8.874°W); 2. Parque das Virtudes, Porto (41.145°N, 8.619°W); 3. Santo Tirso (41.313°N, 8.446°W); 4. Crestuma (41.069°N, 8.503°W); 5. Póvoa de Lanhoso (41.586°N, 8.281°W); 6. Lindoso/Poço da Gola (41.858°N, 8.223°W); 7. Parada de Ester (40.932°N, 8.059°W); 8. Leonte (41.771°N, 8.150°W); 9. Vila Real (41.301°N, 7.737°W); 10. Pendilhe (40.908°N, 7.837°W); 11. Oliveira do Hospital (40.360°N, 7.893°W); 12. Miradouro dos Piornos (40.319°N, 7.573°W); 13. Sabugal (40.350°N, 7.094°W); 14. Montesinho (41.939°N, 6.765°W); 15. Barrocal do Douro (41.429°N, 6.346°W); 16. Vale do Rossim (40.403°N, 7.586°W); 17. Fafião (41.705°N, 8.089°W); 18. Castro Laboreiro (42.028°N, 8.160°W); 19.

Oia (42.002°N, 8.876°W); 20. Ribadavia (42.297°N, 8.137°W); 21. Setados (42.083°N, 8.388°W).

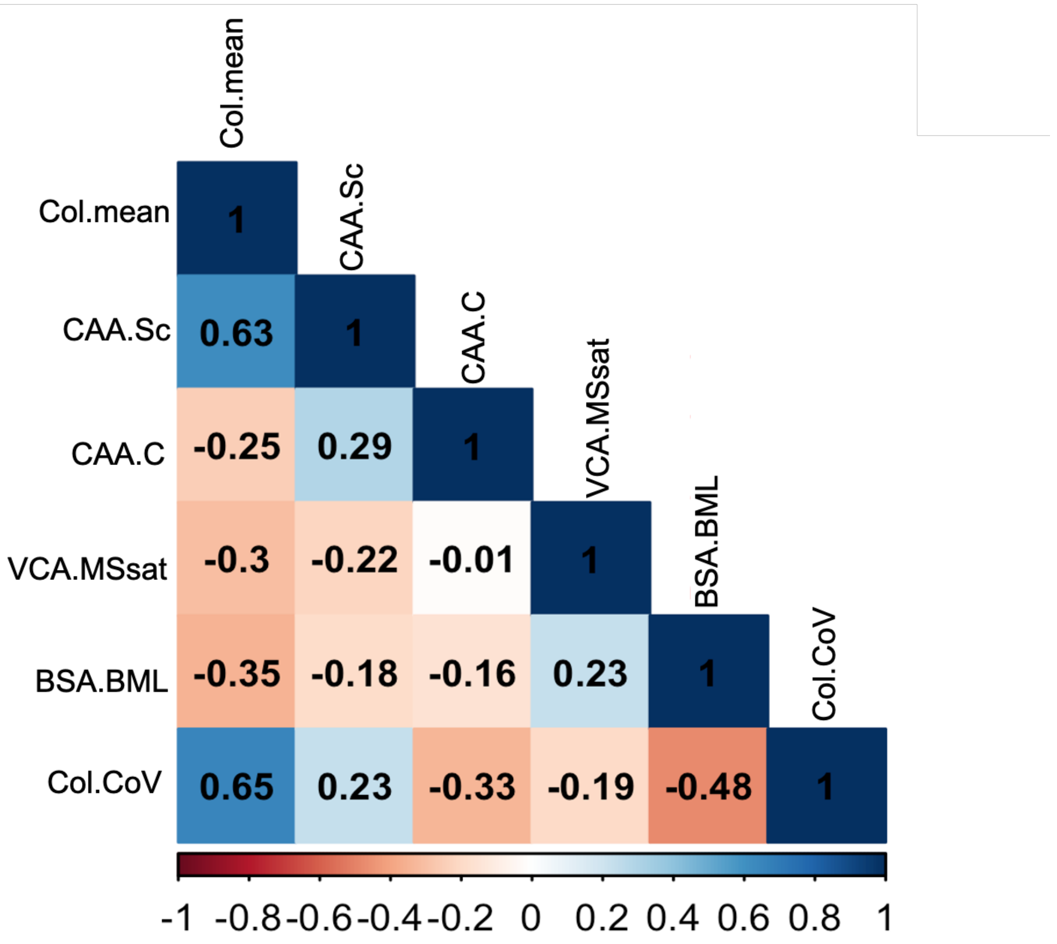


**Figure S2**. Level of correlation between the colour-pattern variables selected from quantitative colour-pattern analysis (QCPA). 1) Col.mean: chromatic edge contrast (mean of the chromatic edge distribution), 2) CAA.Sc : colour diversity (Simpson colour diversity describing how evenly the colour/luminance classes are represented inside a pattern), 3) CAA.C: geometric complexity (geometric complexity of colour patterns based on the number of colour transitions), 4) VCA.MSsat: saturation contrast (weighted mean of pattern RNL saturation contrast, 5) BSA.BML: boundary luminance contrast (average luminance difference of boundaries between colours across the image), 6) Col.CoV: colour variance (Covariance of the chromatic edge distribution). From the extracted original 157 variables from the QCPA pipeline, for variables organised as triplets (horizontal: *.hrz*, vertical: *.vrt*, and combined), we retained only the combined metrics because the three were highly correlated. (See van den Berg et al., 2020) for the complete list of QCPA outputs and further details).

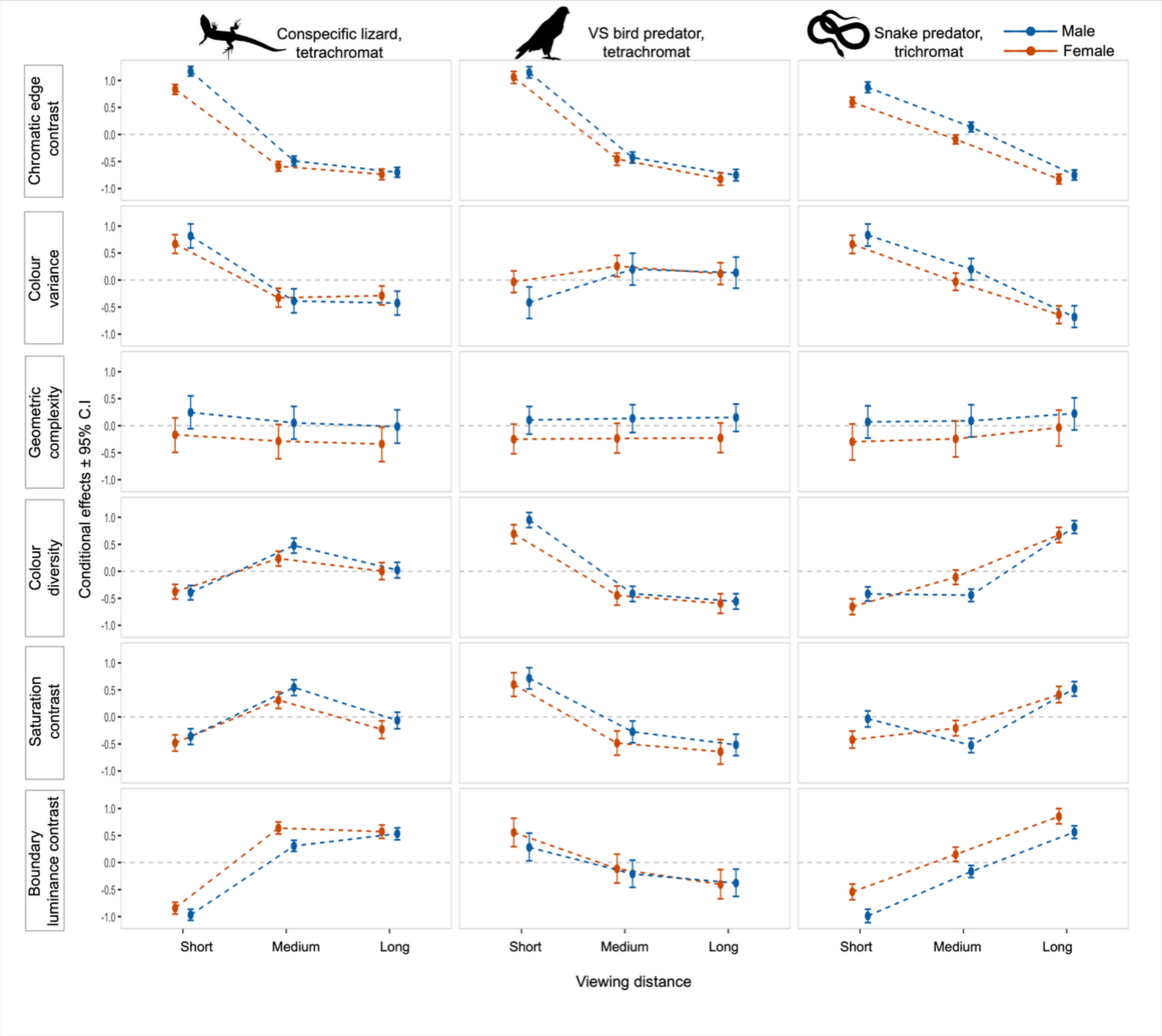


**Figure S3**. Predicted values for six standardised dorsal colour-pattern components of *Podarcis lusitanicus* lizards at short, medium, and long viewing distances, accounting for body size, modelled for three observer visual systems: conspecific lizard, bird predator, and snake predator. Predicted values are shown as conditional effect estimates ± 95% credible intervals (CI). Viewing distances modelled: 1) conspecific lizard: 0.5m (short), 1.5m (medium), 2m (long); 2) Bird predator: 1.5m (short), 3m (medium), 4m (long); (3) snake predator: 1m (short), 1.5m (medium), 3m (long).  Points represent posterior median predictions at each distance level, and error bars indicate 95% credible intervals. The dashed horizontal line at zero indicates the mean of the standardised response, providing a reference for whether predicted values are above or below average. All silhouettes were obtained from

PhyloPic.org. The conspecific lizard silhouette is attributed to Titouan Montessui (licensed under CC BY 4.0, https://creativecommons.org/licenses/by/4.0/), while the violet-sensitive bird and snake silhouettes were sourced from openly licensed images.

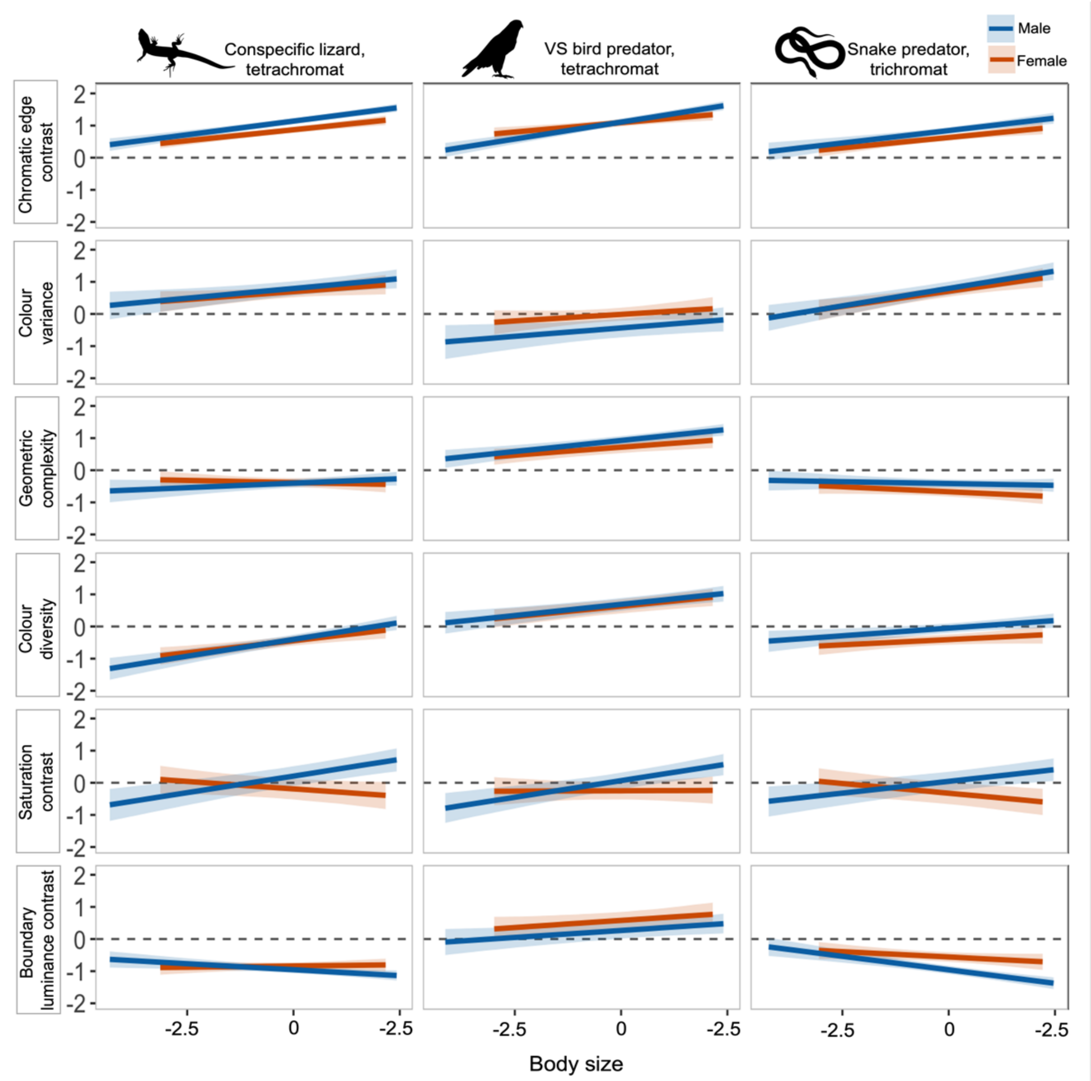


**Figure S4**. Predicted values for six standardised dorsal colour-pattern components of *Podarcis lusitanicus* lizards across a range of body sizes (standardised), accounting for different viewing distances, modelled for three observer visual systems: conspecific lizard, bird predator, and snake predator. Predicted values are shown as conditional effect estimates ± 95% credible intervals (CI). Viewing distances modelled: 1) conspecific lizard: 0.5m (short), 1.5m (medium), 2m (long); 2) Bird predator: 1.5m (short), 3m (medium), 4m (long); (3) snake predator: 1m (short), 1.5m (medium), 3m (long). Lines represent posterior median predictions across the body size gradient; with 95% credible intervals. The dashed horizontal line at zero indicates the mean of the standardised response, providing a reference for whether predicted values are above or below average. All silhouettes were obtained from

PhyloPic.org. The conspecific lizard silhouette is attributed to Titouan Montessui (licensed under CC BY 4.0, https://creativecommons.org/licenses/by/4.0/), while the violet-sensitive bird and snake silhouettes were sourced from openly licensed images.
